# Sub-2 Å Ewald Curvature Corrected Single-Particle Cryo-EM

**DOI:** 10.1101/305599

**Authors:** Yong Zi Tan, Sriram Aiyer, Mario Mietzsch, Joshua A. Hull, Robert McKenna, Joshua Grieger, R. Jude Samulski, Timothy S. Baker, Mavis Agbandje-McKenna, Dmitry Lyumkis

**Author notes:** These authors contributed equally to the work (Yong Zi Tan, Sriram Aiyer, Mario Mietzsch). For materials and correspondence*: Mavis Agbandje-McKenna and Dmitry Lyumkis.

## Abstract

Single-particle cryogenic electron microscopy (cryo-EM) provides a powerful methodology for structural biologists, but the resolutions typically attained with experimentally determined structures have lagged behind microscope capabilities. Here, we have exploited several technical solutions to improve resolution, including sub-Angstrom pixelation, per-particle CTF refinement, and most notably a correction for Ewald sphere curvature. The application of these methods on micrographs recorded on a base model Titan Krios enabled structure determination at ∼1.86-Å resolution of an adeno-associated virus serotype 2 variant (AAV2), an important gene-delivery vehicle.

---

Single-particle cryo-EM has become a powerful tool for macromolecular structure determination, owing largely to numerous technical advances over the past decade^1^. Whereas near-atomic resolution (~3–4 Å) can now be obtained almost routinely, achieving resolutions below ~2.5 Å remains challenging, and only one experimental cryo-EM structure has broken the nominal 2 Å barrier^2^. We sought to address a number of factors limiting the resolution of structure determination by single-particle cryo-EM. In these analyses, we studied a variant of adeno-associated virus (AAV) serotype 2 containing a single amino-acid substitution, L336C. The AAV2_L336C_ variant is of particular biological interest, as it is defective in genome packaging and is associated with reduced infectivity^3, 4^ AAVs are single-stranded DNA viruses that infect vertebrates^5^ and are thereby attractive vehicles for gene delivery^5, 6^, with AAV2 being one of the most popular serotypes for such applications. The AAV viral capsid is formed by an icosahedral (T=1)^5^ arrangement of 60 viral protein (VP) monomers, and has a molecular weight of ~3.9 MDa and a shell diameter of ~250 Å. The three related capsid proteins, VP1, VP2, and VP3, share a common core sequence and occur in a predicted 1:1:10 ratio^7^. AAV was particularly suited to our cryo-EM studies because: 1) it is relatively small for a virus and can be packed across cryo-EM grid holes in reasonably thin ice; 2) it can be stably assembled into homogeneous virus-like particles (VLPs) devoid of genomic material; and 3) it has icosahedral symmetry, which increases the number of asymmetric subunits in the dataset by 60-fold for each particle imaged.

To image AAV2_L336C_ particles, we used a base model Titan Krios operating at 300 keV with a K2 summit detector, without the use of newer technologies such as phase plates^8^, Cs correctors^9^, and energy filters^2^. For data collection, parameters such as aperture choice, beam size, magnification, camera settings, choice of stage shift for targeting and defocus range build upon optimal conditions elucidated in previous high-resolution single-particle collections^2, 10^ and are elaborated in supplementary note 1. After recording a Zemlin tableau from a gold-coated cross-grating replica calibration grid, which revealed a coma-free aligned beam and evidence for 1.44 Å gold diffraction spots (Supplementary Fig. 1), we proceeded with collecting cryo-EM micrographs of AAV2_L336C_ (Supplementary Fig. 2). For data processing, multiple procedures resulted in statistically significant improvements in resolution, as evidenced by changes across most frequency ranges within Fourier Shell Correlation (FSC) curves, summarized in Figure 1a. Improvements are described in spatial frequency shells, since at higher resolutions, statistically significant gains are characterized by incrementally smaller increases in nominal resolution values. First, we removed particle images with the greatest angular uncertainty based on conventional scoring criteria in either Relion^11^ or Frealign^12^ and adjusted the weights for how different particles contribute to the reconstruction. We then performed per-particle CTF estimation using GCTF^13^ and subsequently refined these values in *cis*TEM^14^, providing a cumulative gain of 46 resolution shells (~0.3 Å). Correcting for magnification anisotropy (estimated at ~1%)^15^ provided gains in resolution by 35 shells (0.21 Å). Notably, map resolution increased by 15 shells (0.09 Å) after correcting for the curvature of the Ewald sphere^16^, which has been predicted, but not previously demonstrated with experimental single-particle cryo-EM data (discussed further below). Additionally, per-frame reconstructions allowed us to determine which of the 70 frames contained the most information content. Reconstructions from individual frames (each receiving a dose of 0.32 e^−^/Å^2^), provided maps with resolutions ranging between 2.1 to 3.4 Å (Supplementary Fig. 3). Discarding the first 4 frames, which contained the largest beam-induced movement, improved the map by 9 resolution shells (0.05 Å), and frames 5–19 could also be combined to produce a largely identical reconstruction to one composed from frames 5–70 (Fig. 1a and Supplementary Fig. 4). Finally, we found that correcting for the rotational particle movement through the course of the movie by refining the orientations of groups of five-frame averages improved low spatial frequency FSC values and the quality of the map, although the nominal value remained largely unchanged. Cumulatively, the above procedures resulted in a total gain of 71 resolution shells (~0.4 Å). As previously demonstrated^17^, the summation of individual gains is not equal to the cumulative improvement, as the effects are not additive.

**Figure 1.**
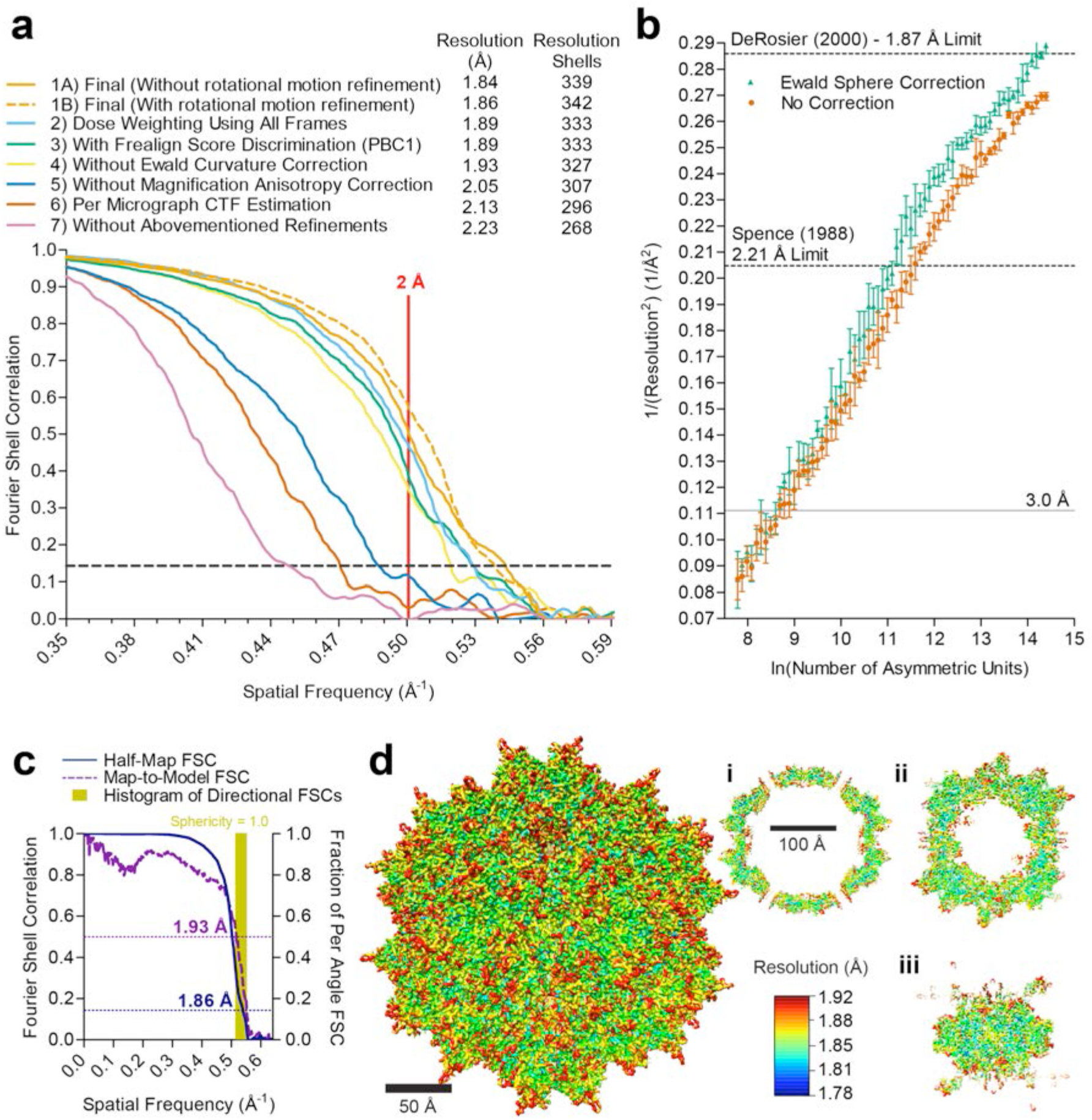
Procedures and implications for obtaining a sub-2 Å resolution reconstruction of AAV2_L336c_. (**a**) Fourier shell correlation (FSC) curves showing independent contributions of each operation to the final resolution. Each FSC curve was generated by “turning off’ one of the respective operations from the final refinement (1A). Correcting for rotational beam-induced movement (1B) improves the FSC at low spatial frequency. Discriminating particles based on their score (3) led to a lower resolution reconstruction (1.89 Å) as compared to equally weighting all particles (1.84 Å). Cumulative loss of “turning off’ all the tested operations is indicated by (7). (**b**) ResLog^26^ plot of the dataset, with (green) and without (red) Ewald sphere curvature correction in Frealign9. Five replicates were performed for each data point with random particle distributions. The resolution limits predicted by DeRosier^19^ and Spence^21^ for a 250 Å macromolecule are indicated. (**c**) Fourier shell correlation (FSC) curves describing the half-map (blue) and map-to-model (purple) resolutions, as well as histogram of directional resolutions sampled evenly over the 3DFSC^27^ (yellow), and corresponding sphericity value. (**d**) 1.86 Å reconstruction of the AAV2 viral capsid using single-particle cryo-EM, colored by local resolution^28^. Slices through the reconstruction displayed in sub-panels i-iii start from the capsid center (i) and move progressively towards the front (ii and iii). These show few deviations from the global resolution, with the best local resolution at 1.78 Å (primarily located inside the core region of the capsid shell).

The above results revealed that correcting for the curvature of the Ewald sphere is pertinent to experimental reconstructions in high-resolution single-particle cryo-EM analysis, warranting further investigation. Most 3D reconstruction algorithms assume that images correspond to direct projections of the 3D object, in accordance with the central slice projection theorem^18^. However, several aspects of cryo-EM data acquisition invalidate this approximation at resolutions approaching true atomic^19, 20^. Most notably, imaged objects have finite thickness along the optical axis of the microscope, which results in an inherent focus gradient during imaging. The focus gradient alters the phases and amplitudes associated with each Fourier coefficient, and the effects become progressively more pronounced at higher resolutions, lower accelerating voltages, or for thicker specimens^21^. Various schemes to estimate and correct for the curvature of the Ewald sphere have been developed^16, 19, 20, 22^, but no experimental reconstruction from a single-particle macromolecular sample has to date demonstrated improvements from taking Ewald curvature into account. The “simple-insertion” method for Ewald curvature correction implemented in Frealign9 will insert the data for each particle image twice into correct Fourier coefficients related by Friedel symmetry^16^. This procedure, performed during reconstruction, resulted in an increase of 15 resolution shells within our final map (Fig. 1a). We then evaluated the effect of the correction at lower resolution by reducing the number of particles in the reconstruction. Randomly selected subsets of the data containing an approximately equal defocus range were used to perform reconstructions with incrementally smaller numbers of particles. As few as ~60 particles (3,600 asymmetric units) were sufficient to produce a ~3.5 Å map, whereas ~120 particles (7,200 asymmetric units), and all larger subsets, were sufficient for <3 Å reconstructions (Supplementary Fig. 5). These maps could be used to evaluate Ewald curvature effects as a function of resolution for the ~250 Å diameter particle. Noticeable gains appeared at ~2.4–2.3 Å, and a final improvement of 15 shells (~0.1 Å) for the best reconstruction (Fig. 1b). The gains follow an increasing trend at higher resolution, as the effects of Ewald curvature become more pronounced at higher electron scattering angles^19^. Furthermore, the correct handedness of a reconstruction can be explicitly determined when accounting for the effects of the Ewald sphere^16^. Specifically, the handedness of the reconstruction defines how the Fourier coefficients are substituted in the reconstruction and whether an inversion operation must be applied to the Fourier coefficients (Supplementary Fig. 6).

The final map of AAV2_L336C_ had a global resolution of 1.86 Å, with a largely homogeneous local resolution distribution within the core of the capsid shell that drops to ≥1.92 Å at the solvent exposed surfaces (Fig. 1c-d and Supplementary Table 1). Using this map, an atomic model was derived for the common region of the VP monomer, residues 226 to 735 (VP1 numbering), which was symmetry expanded by icosahedral matrix multiplication to produce the full 60-mer viral capsid. As in previously reported AAV structures, the VP1u, VP1/2 common sequence, and the N-terminus of common VP3 are disordered. The final model corresponded closely to the map, with good statistics (Supplementary Table 1), including a high EM-Ringer^23^ score of 8.49 and a correlation coefficient following model refinement in Phenix of 0.849^24^. The EM-Ringer score reflects accuracy of fit between model and map based on side-chain rotameric positions. At this resolution, the map is of sufficient quality to see numerous features with unprecedented detail (Fig. 2 and Supplemental Movie 1) including: 1) the backbone tracing with well-defined carbonyls; 2) explicit structure to most side-chains, rotamers, holes in aromatic residues, as well as prolines and associated puckers; 3) ordered solvent throughout the structure, including primary and secondary hydration shells; and 4) the distinct appearance of density for individual oxygens of carboxylate groups, which occasionally begin to show traces of H-bonding geometry.

**Figure 2.**
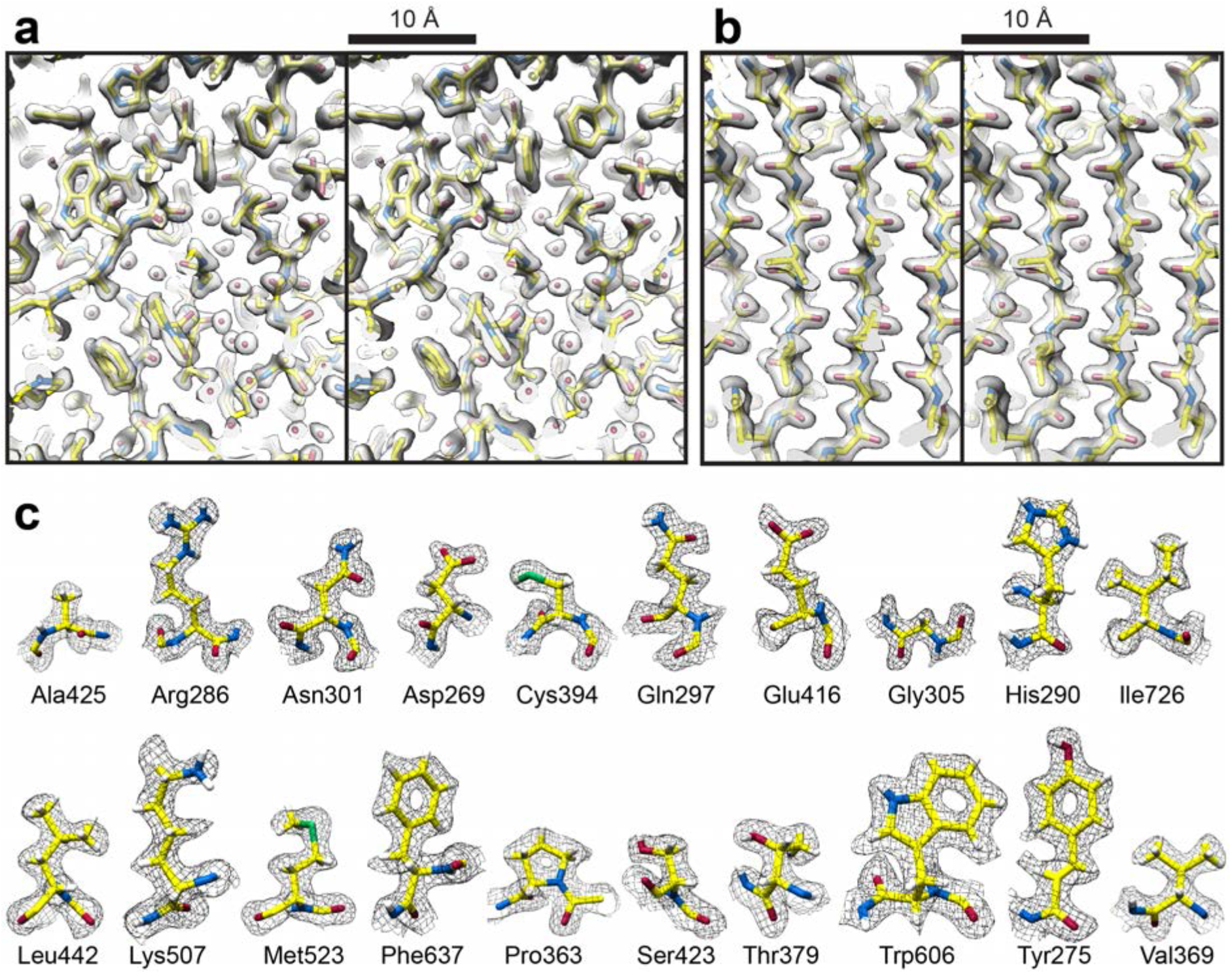
High-resolution information within the 1.86 Å reconstruction of AAV2_L336C_. (**a**) Stereo view of a slice through the map and model containing both amino acid residues and water molecules. (**b**) Stereo view of a slice through a beta sheet. (**c**) Map densities for each of the 20 types of amino acid residues. The amino acid residues are shown as stick representation and colored according to atom type: C=yellow, O=red, N=blue, S=green inside either a translucent solid density (**a,b**) or black mesh density map (**c**). H=white atoms are displayed for (**c**).

Efforts to improve the AAV gene delivery system have focused on structure-function analysis of the viral life cycle and of engineered capsids to improve therapeutic efficacy. The 1.86-Å resolution structure of AAV2_L336c_ represents the most accurately interpreted AAV capsid model thus far (Fig. 3). This particular variant also exhibits specific structural changes that are clearly captured in the density map, and these changes may be associated with infectivity defects (Supplementary Note 2 and Supplementary Fig. 7). High-resolution structural information can aid the annotation of: 1) water networks required to stabilize the capsid structure assembly and involved in its function; 2) the protonation states of acidic and histidine residues important for interactions in the endo/lysosomal pathway; 3) capsid interactions with the transcription machinery and during capsid assembly; and 4) precise receptor and antibody interactions. Details from such analyses can guide the engineering of AAVs at specific residues to eliminate interactions, such as those with pre-existing host immune system molecules, or improve function, such as specific tissue targeting. Notably, the number of known sites of interaction between the AAV ligand and host receptor/antibody far exceeds the number of experimentally derived AAV models. For this reason, the fact that high-resolution structures of AAV variants can be derived with as few as ~100 particles (Supplementary Fig. 5) is noteworthy and will accelerate the compilation of a comprehensive structural understanding of AAV:host interactions^5^.

**Figure 3.**
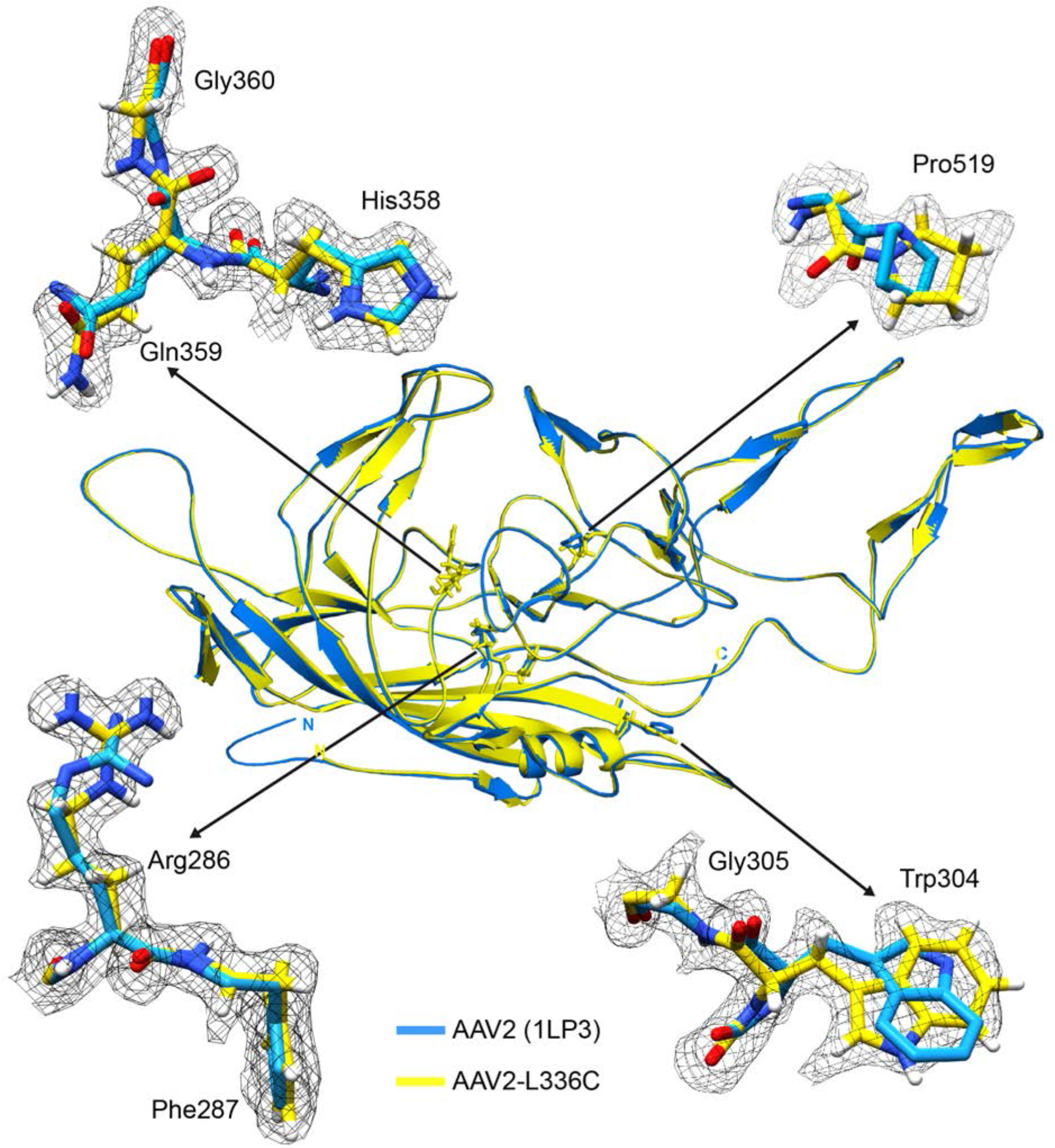
High-resolution map aids in placement of correct rotamers for AAV2_L336C_. At 1.86 Å (map in black mesh), uncertainty in rotamer and backbone conformation in a previous model of AAV2 (1LP3, in cyan) determined by X-ray crystallography to 3.0 Å resolution can now be accurately modelled (yellow).

The methods described herein provide a feasible route toward true atomic resolution in cryo-EM single-particle analysis. While we used a well-behaved sample for this work, the modest amount of time (3.5 days collection) and equipment (a base model Titan Krios and K2 Summit camera) used for this reconstruction would make our strategy generally applicable, even though the relative gains will differ by specimen (Supplementary Fig. 8). Correcting for the curvature of the Ewald sphere should be incorporated into reconstruction algorithms and may have particular relevance for improving resolution for samples collected at lower microscope accelerating voltage^25^. Finally, our sub-2 Å resolution reconstruction AAV2_L336C_ also provides new insights into AAV life cycle and biology that will be invaluable for improving the effectiveness of AAV as a delivery vehicle in gene therapy applications.

## Acknowledgements

Molecular graphics and analyses were performed with the UCSF Chimera package (supported by NIH P41 GM103331). We thank Bill Anderson at TSRI for help with EM data collection. We also thank David DeRosier and Gordon Louie for critical reading of the manuscript. The work was supported by Agency for Science, Technology and Research Singapore (to Y.Z.T.); NIH R01 GM109524 (to R.M. and M.A.M.), R01GM033050 (to T.S.B.), and NIH R01AI136680-01 (to D.L.). We acknowledge consultations with Bridget Carragher and Clinton S. Potter. Some of the work was performed at the National Resource for Automated Molecular Microscopy at the Simons Electron Microscopy Center which is supported by National Institute of General Medical Sciences (GM103310), Simons Foundation (SF349247) and NYSTAR.

## Accession codes and deposition

All raw movie frames, micrographs, the particle stack and relevant metadata files will be deposited into EMPIAR. The electron density map will be deposited into EMDB. The model will be deposited into PDB.

## Author Contributions

J.G. generated the baculovirus construct. J.A.H. purified the sample. D.L. and S.A. vitrified the sample and collected the data. Y.Z.T., D.L., and S.A. processed the data. D.L., M.M., Y.Z.T. and S.A. built and refined the model. M.A.M., R.J.S., T.S.B. and R.M. conceived the variant study. D.L. and Y.Z.T. conceived the high-resolution study. T.S.B., M.A.M. and D.L. supervised throughout the experiment. All authors read and contributed to the manuscript.

## Supplementary Note 1 | Imaging Conditions

For sample preparation, we used gold grids to reduce beam-induced movement^29^. For microscopy, we selected the smallest available C2 condenser aperture (70 μm on our Titan Krios) to maximize beam coherence. The spot size was selected to yield a beam diameter of ~2 μm to allow for complete illumination of the imaged hole, while ensuring that the imaged area is within range of parallel illumination^30^. During data acquisition, we used a relatively high magnification (37kx, corresponding to a pixel size of 0.788 Å) to boost the detective quantum efficiency (DQE) of the direct electron detector (Gatan K2 Summit) at a fixed spatial resolution^31, 32^. Super-resolution mode boosted the final pixel size to 0.394 Å, which facilitates reaching close to or even beyond physical Nyquist in the reconstructed map^31^ while maintaining an otherwise identical field of view. During imaging, the microscope stage shift^10^ (rather than beam tilt-induced image shift) was used to center the specimen field of view so as to minimize beam tilt (and the concomitant effects of coma) within the final images, and a 60 s delay before the exposure was implemented to reduce grid movement due to thermal instability. A frame rate of 20 frames / sec (50 ms frames) was used to correct for beam-induced motion^33^, although a running average of 3 sequential frames was subsequently determined to be an optimal setting for frame alignment. A dose of ~4 e^−^/pixel/s on the K2 Summit detector was selected to reduce coincidence loss and maximize the camera DQE^31^. With the above procedures, a coma-free microscope alignment allowed for observation of the 2.36, 2.04 and 1.44 Å diffraction spots from the gold-shadowed cross-grating replica calibration grid (Supplementary Fig. 1). This result indicated that the aligned microscope was capable of capturing high resolution information in recorded images. While the abovementioned settings may differ slightly between microscopes and facilities, any employed procedures should result in high-resolution diffraction spots, as shown in Supplementary Fig. 1, to maximize data quality. Data collection throughput was aided by automation using Leginon^34^ across the 3.5-day-long session. We collected 1,317 micrographs of AAV2_L336C_ particles, which after extensive pruning and classification (see Methods) provided a dataset of 30,515 particles for refinement (Supplementary Fig. 2 and Supplementary Table 1).

## Supplementary Note 2 | Structural Comparison of AAV2_L336C_ and AAV2_WT_

The density for C336 is clearly ordered in the AAV2_L336C_ map, and the model is considerably improved compared with the prior structure of AAV2_WT_ (Fig. 3)^35^. There is a 1.4 Å shift of the main-chain of C336 and neighboring residues, compared to AAV2_WT_ (Supplementary Fig. 7a). This results in a 0.8 Å widening at the base of the 5-fold channel formed by five symmetry related DE loops (the loop between the βD and βE strands). In addition, the AAV2_WT_ structure is ordered from residue 217 to 735, with the additional N-terminal residues compared to AAV2_L336C_ (residues 226 to 735) occupying the base of the interior opening of the 5-fold channel (Supplementary Fig. 7b). The AAV2_L336C_ variant displays a 23-fold defect in genome packaging compared to AAV2_WT_ and lacks PLA2 activity resulting in a defect in infectivity^3, 4^ This defect was proposed to be due to the inability to expose the PLA2 and potential structural differences to AAV2_WT_. The annotated differences in AAV2_L336C_ support these possibilities. An altered location of the PLA2 domain due to the N-terminal disorder would abrogate its externalization via the 5-fold pore and thus its function.

**Supplementary Figure 1.**
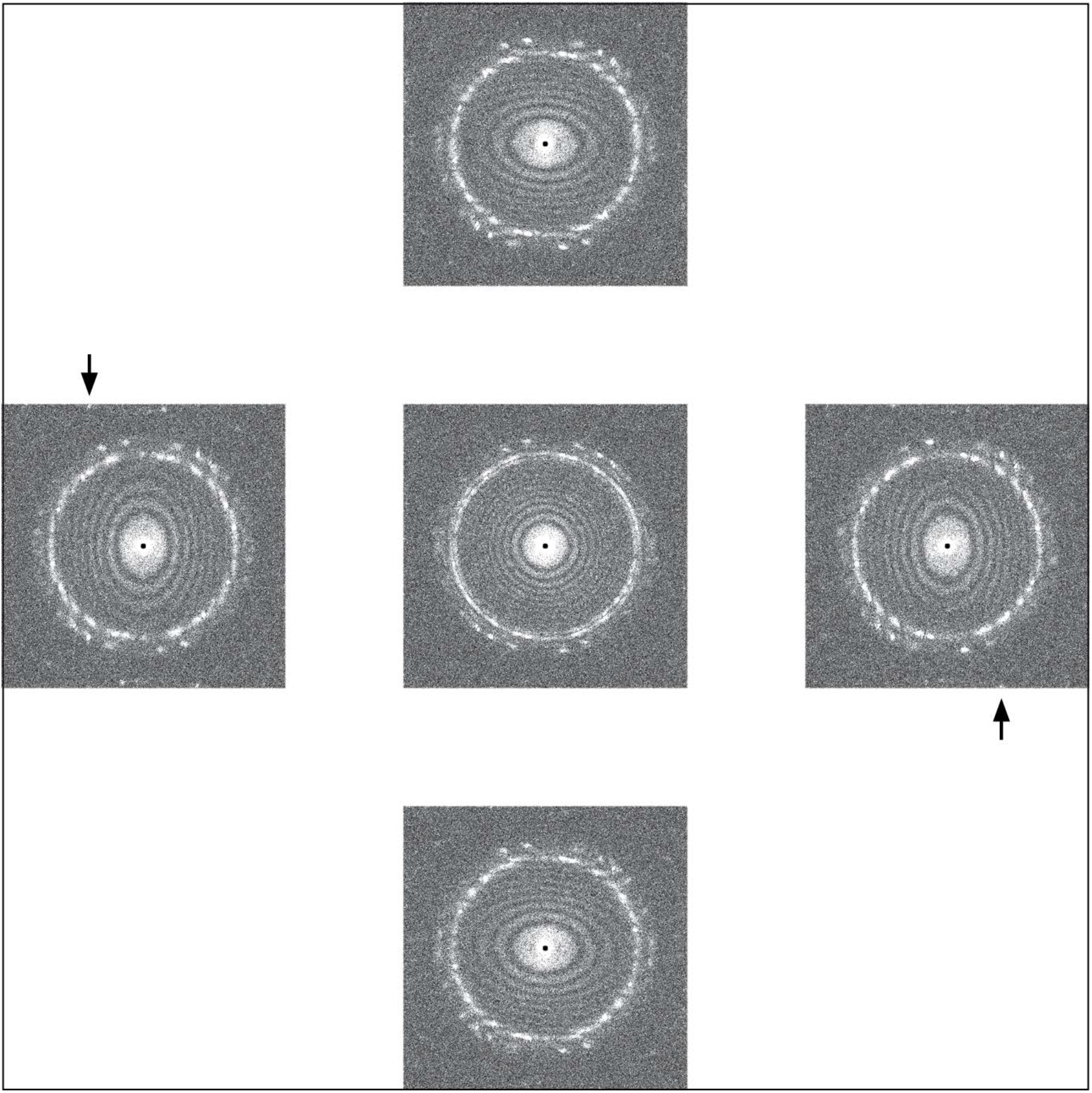
High-resolution information is present within micrographs. Zemlin tableau recorded under cryogenic conditions showing power spectra of images of a gold-coated carbon cross-grating replica grid, acquired with a 5 mrad beam tilt in the respective direction, and subsequent to coma-free microscope alignment. Gold diffraction spots at 2.355 Å (averaging to a nearly concentric ring), 2.039 Å, and 1.442 Å are evident within the series of power spectra. Arrows point to faint diffraction spots visible at 1.442 Å.

**Supplementary Figure 2.**
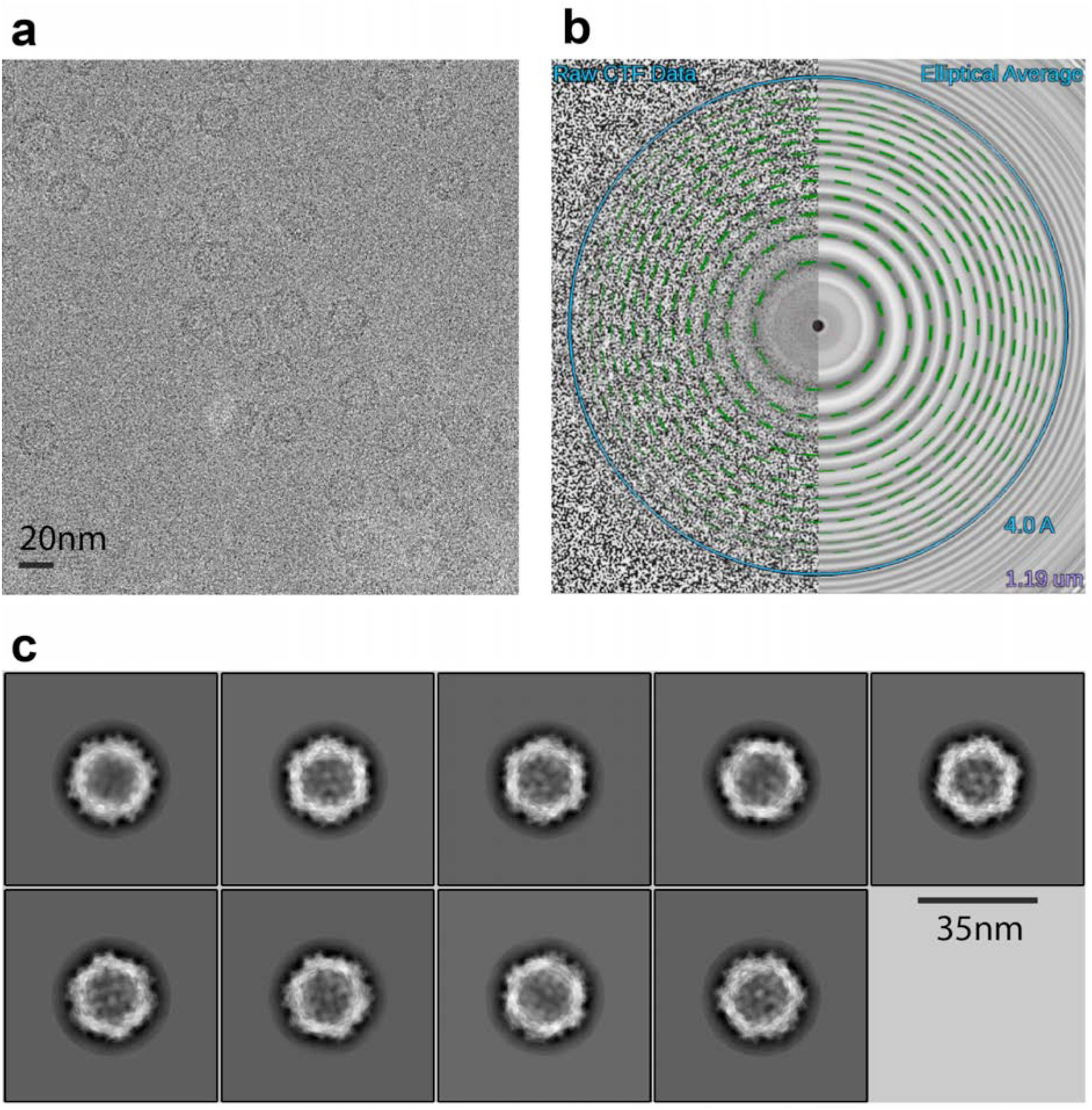
Raw cryo-EM data. (**a**) A cryo-EM micrograph of the AAV2_L336c_ imaged on a Titan Krios with a Gatan K2 summit direct detector at −1.19 μm defocus, estimated using CTFFind4. (**b**) CTF estimation profile from Appion^36^, showing agreement of the Thon rings and estimated CTF profile at 80% confidence to 3.21 Å. (**c**) 2D class averages of AAV2_L336C_ aligned using Relion.

**Supplementary Figure 3.**
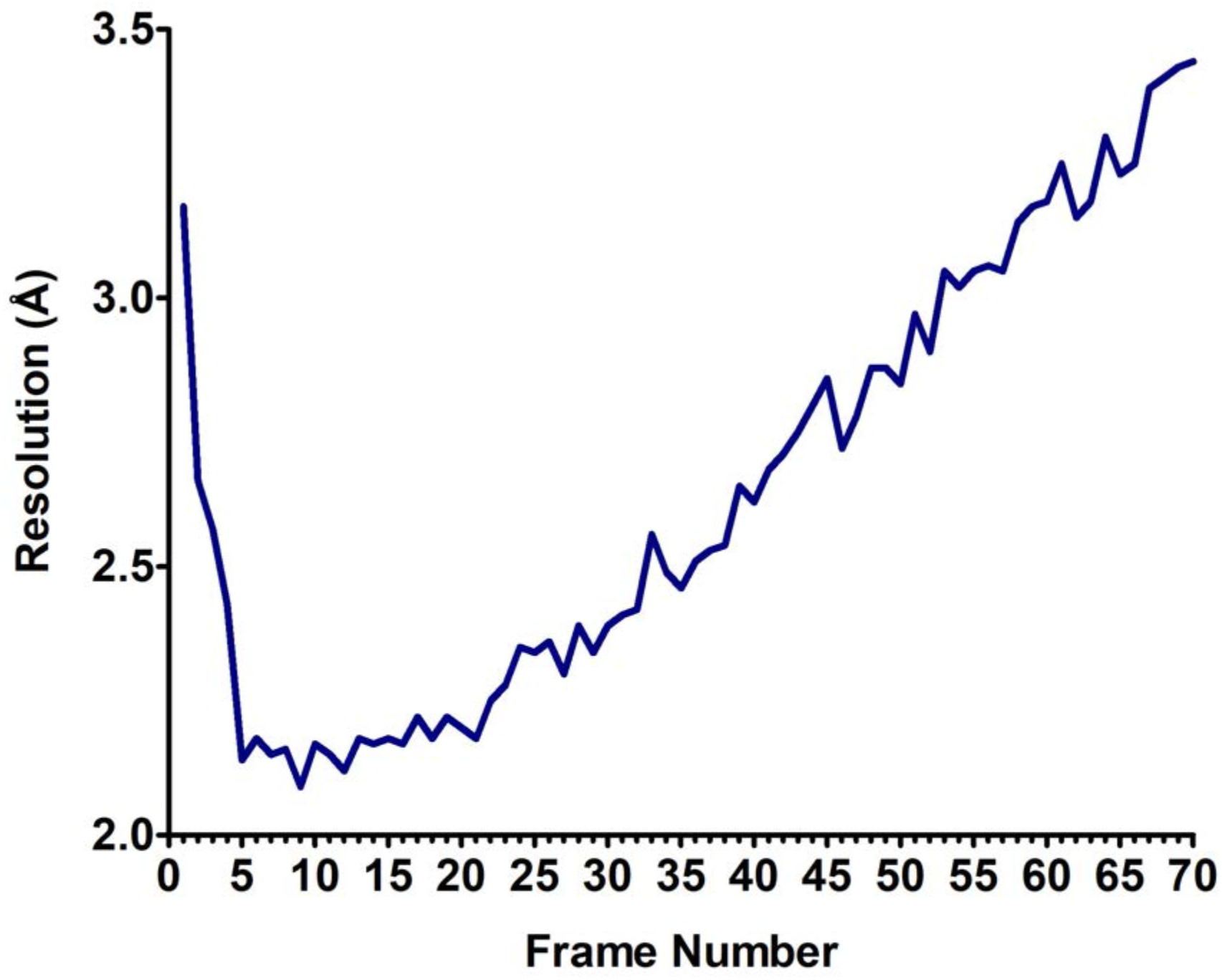
Resolution of individual frame reconstructions. Using the best Euler angles and shifts, reconstructions were computed separately for each of the 70 frames. The resulting resolution shows two trends: the first 4 frames (3.17–2.43 Å) suffered from the initial effects of beam-induced motion; after frame 22, the resolution gradually worsens owing to the cumulative effects of radiation damage.

**Supplementary Figure 4.**
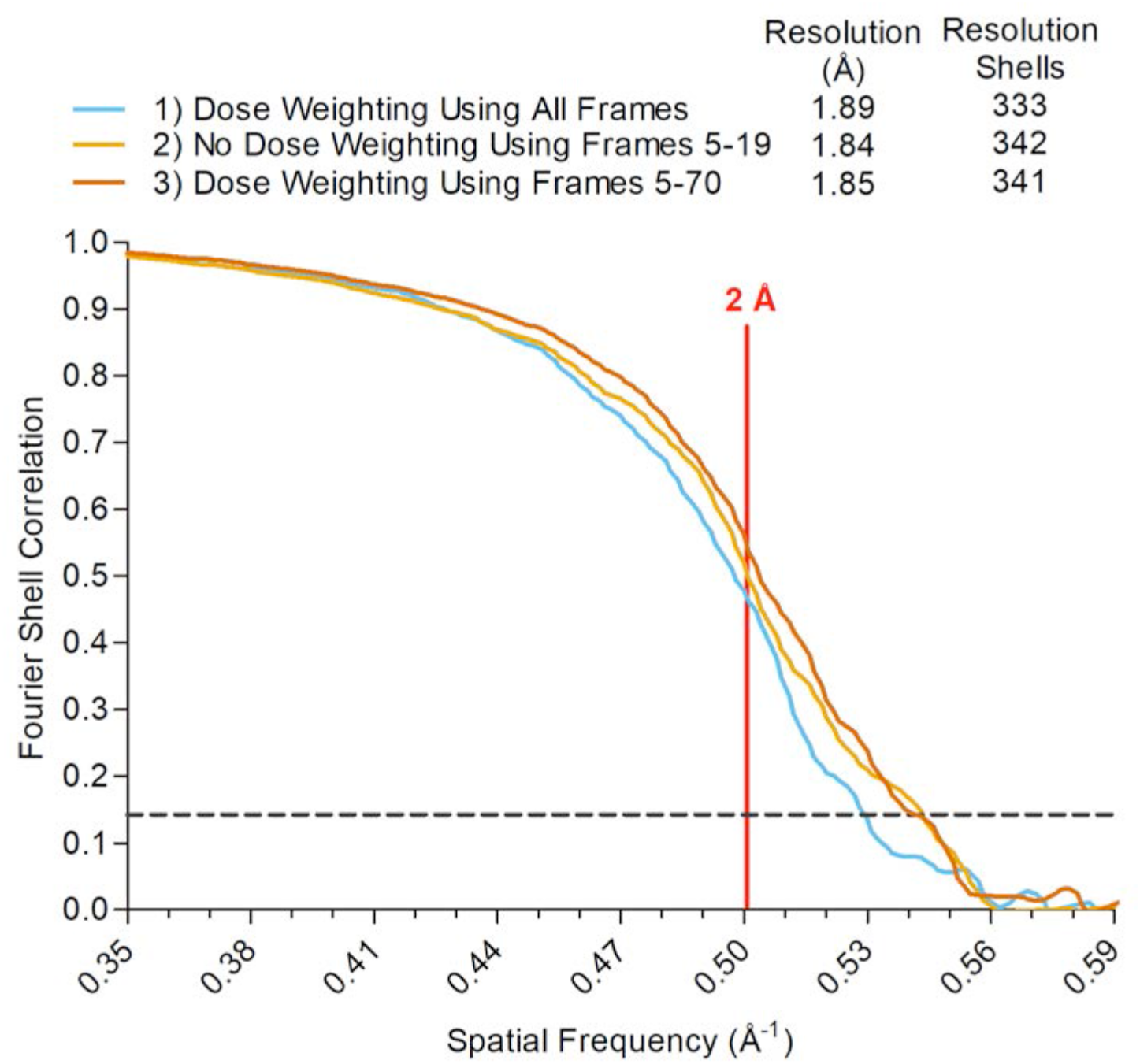
Effect of removal of first frames. Removing the first 4 frames that exhibit the most beam-induced movement, and consequently the lowest resolution (see Supplementary Fig. 3), either from the subset of frames 5–19 (2) or from the cumulative, dose-weighted sum containing frames 5–70 (3) produces measurable resolution gains.

**Supplementary Figure 5.**
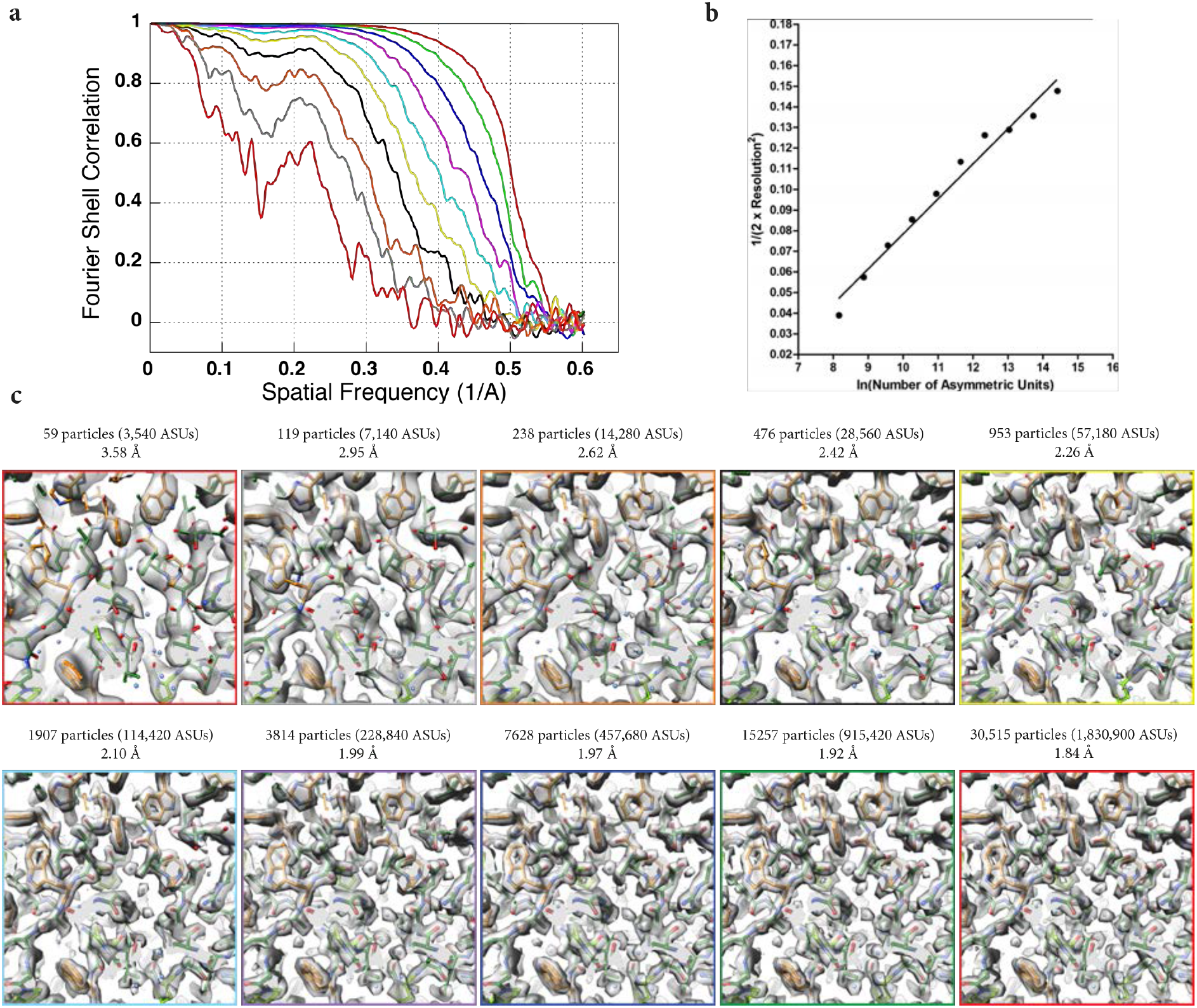
Improvement in resolution with increasing particle number. Reconstructions were performed with subsets of the particles, before rotational motion correction. (**a**) FSC curves, (**b**) a plot of nominal resolution value (expressed in inverse Å^2^) as a function of the number of asymmetric units (ASUs), and (**c**) views of the density maps showing structural features for each corresponding reconstruction. Water molecules and holes in aromatic residues become obvious at beyond 2.10 Å.

**Supplementary Figure 6.**
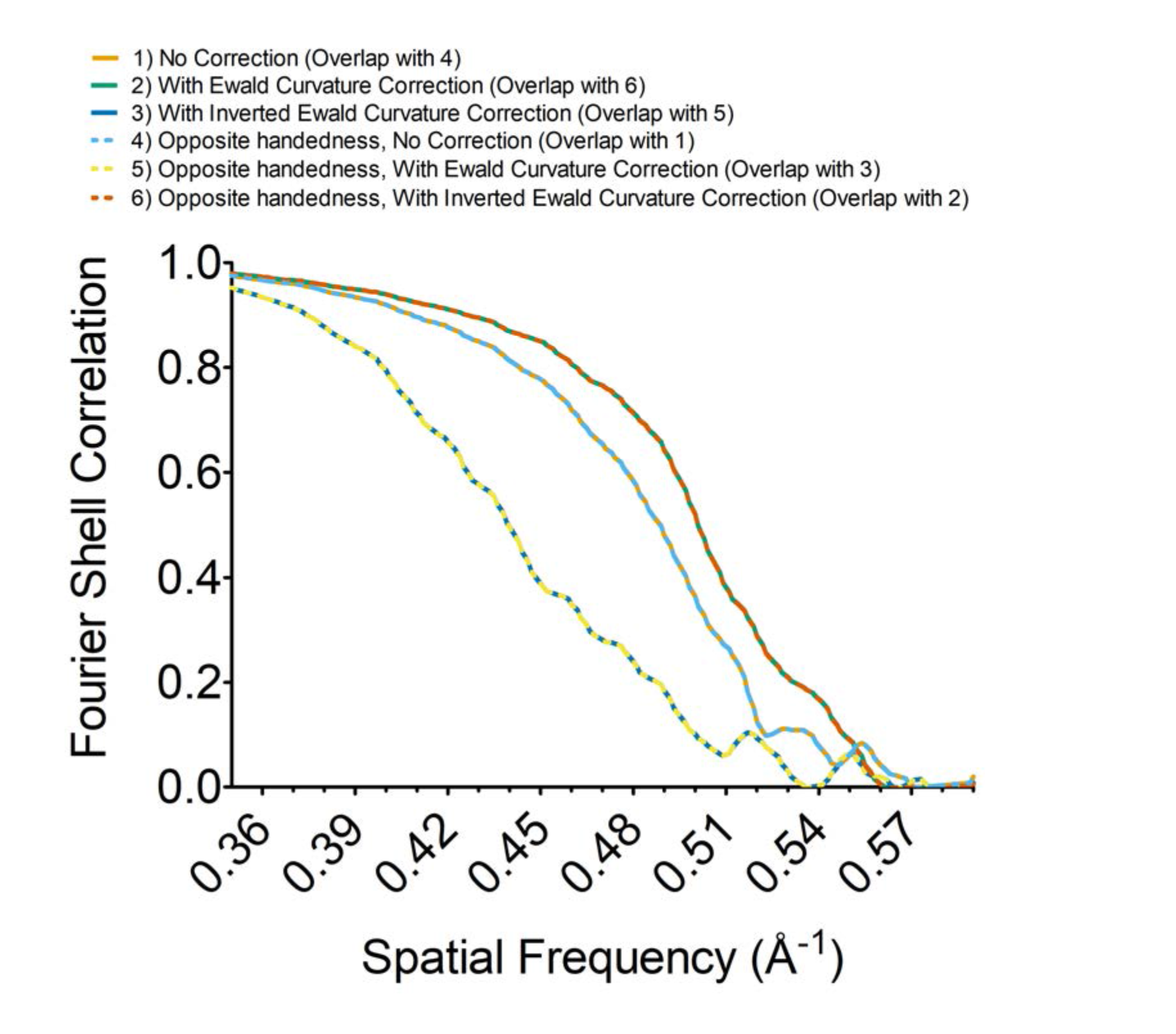
Accounting for Ewald sphere curvature improves cryo-EM resolutions and allows for determination of map handedness. A map reconstructed without correcting for the curvature of the Ewald sphere and using orientation parameters accounting for (1) the correct or (4) opposite map handedness have identical resolutions. Accounting for the curvature of the Ewald sphere during the reconstruction for the correct-handed map improves resolution (2), whereas the same operation performed on a reconstruction of the opposite-handed map decreases the resolution (5). Analogously, an inversion of the Fourier coefficients during Ewald curvature correction, if applied to the correct-handed map, decreases resolution (3), whereas this same operation performed during the reconstruction of the opposite-handed map effectively restores all high-resolution components (6). These sets of operations allow for automated determination of handedness in high resolution cryo-EM maps in the absence of auxiliary data, as previously theorized^16, 22^.

**Supplementary Figure 7.**
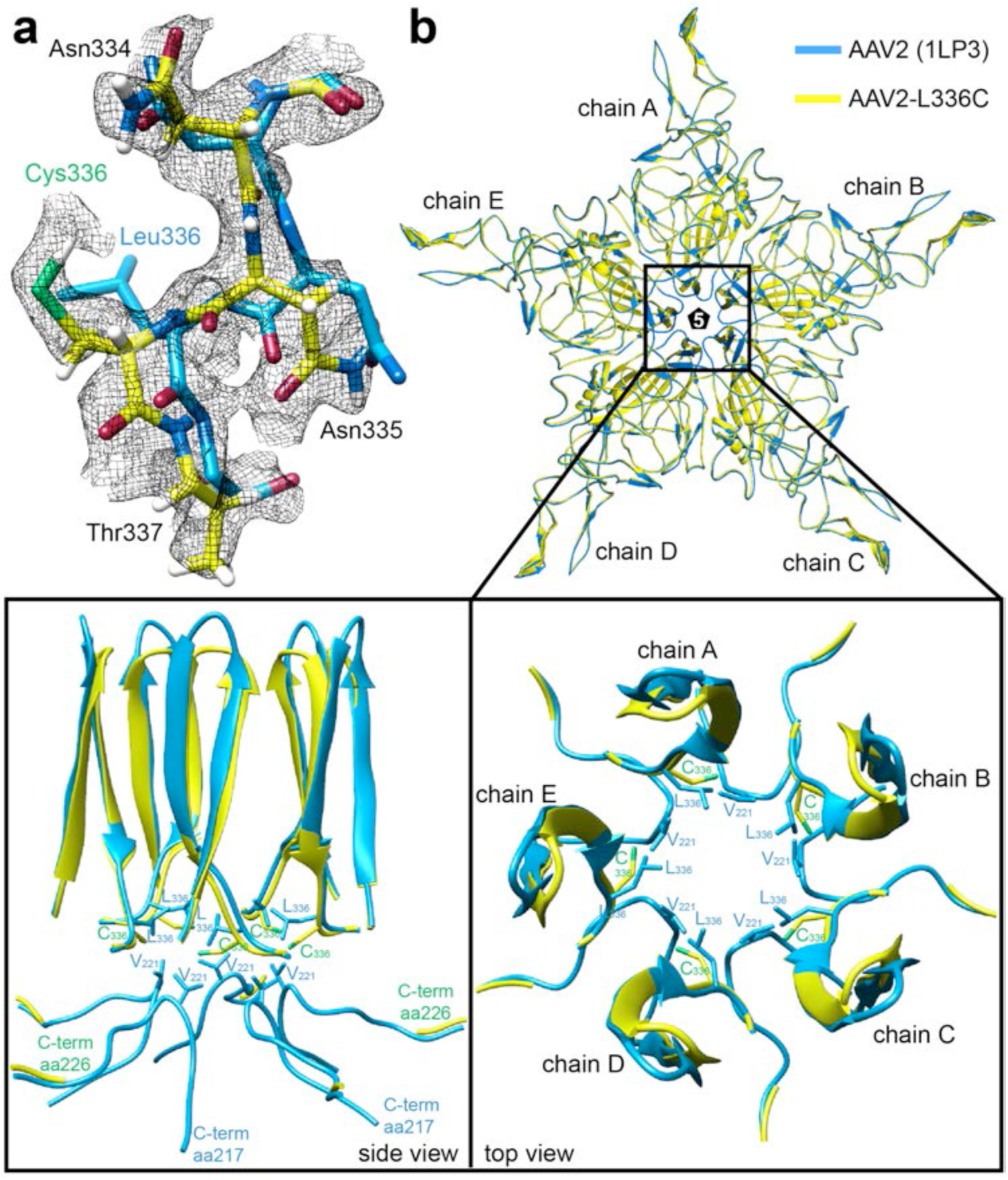
Comparison of AAV2_WT_ and AAV2_L336C_. (**a**) The density map with the modeled residues for the DE-loop for AAV2_WT_ and AAV2_L336C_. The wild type L336 and substituted C336 residues are shown. The AAV2_WT_ has been the highest resolution density map for AAV2 published to date, at 3 Å. The amino acid residues are shown as stick representation and colored according to atom type: C=yellow, O=red, N=blue, H=white, S=green inside a black mesh density map. (**b**) Superposed pentamer models of AAV2_WT_ (blue) and AAV2_L336C_ (yellow). Lower panel show close-up views of the 5-fold region as side view and top view perspectives. Side-chain atoms for individual residues of interest are shown.

**Supplementary Figure 8.**
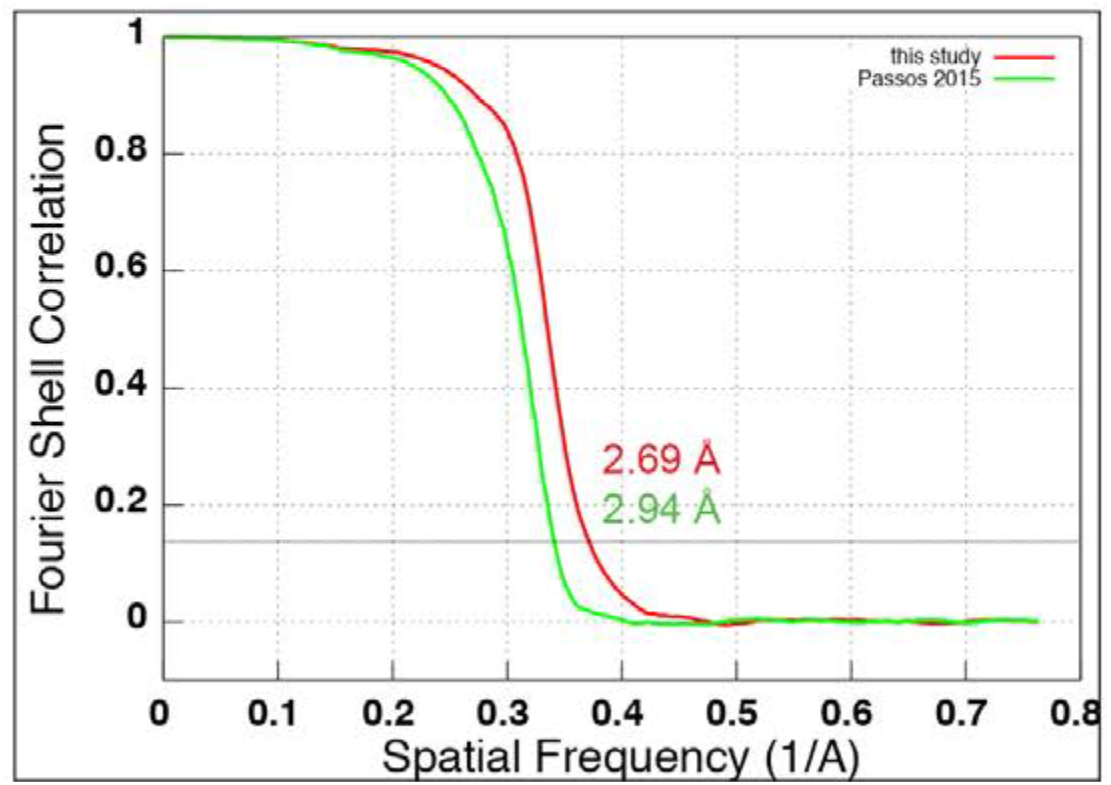
Improvement of a previously published dataset with per-particle CTF estimation. Resolution curves describing reconstructions of the 60S ribosomal subunit from Passos *et al*.^3^ (green) and from this work, after applying per-particle CTF estimation (red). At these resolutions, correcting for the curvature of the Ewald sphere did not significantly improve the map, in line with our observations with AAV2_L336C_.

**Supplementary Table 1.**
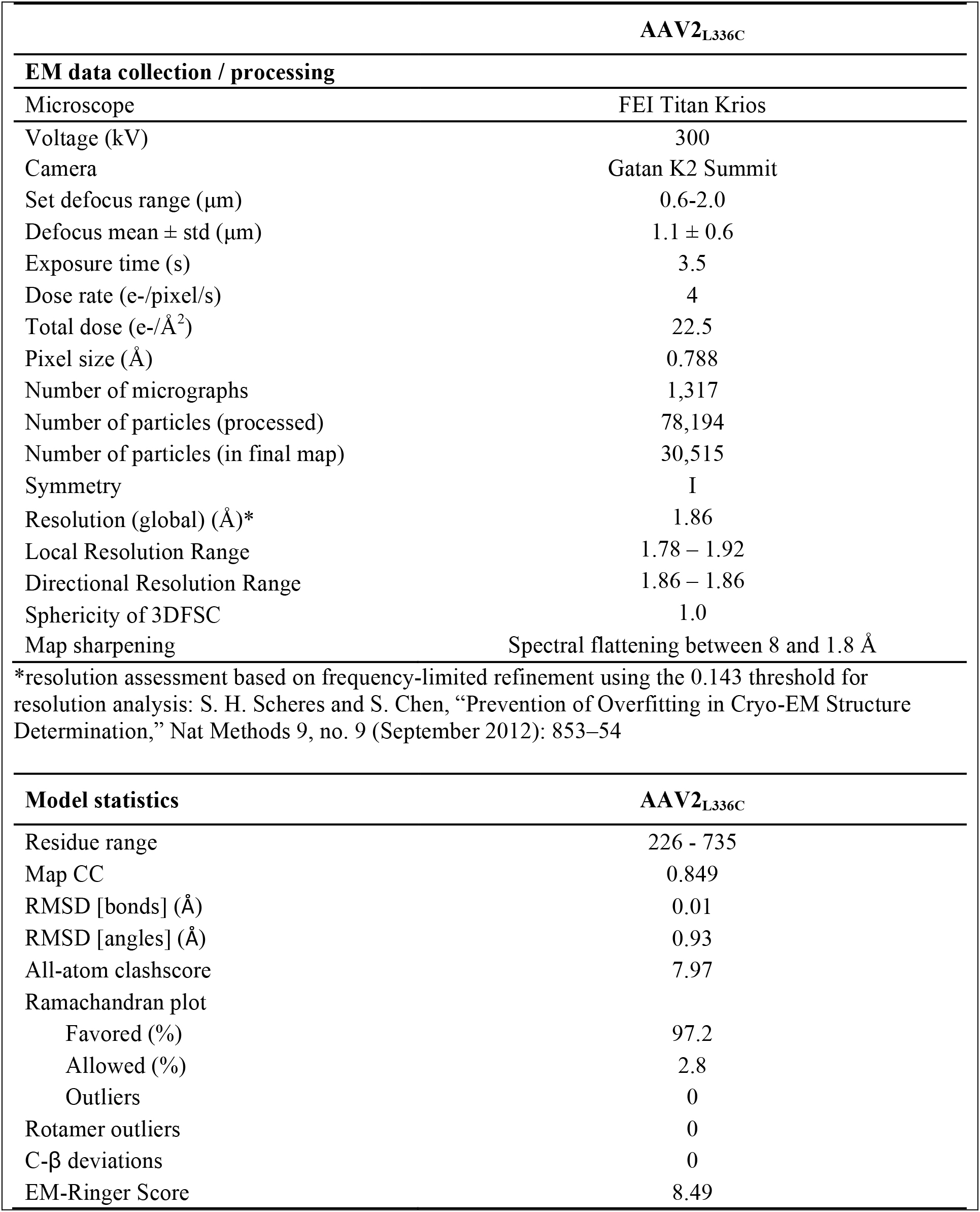
Cryo-EM data collection and modeling statistics.

## Online Methods

### Statistics

For calculations of Fourier shell correlations (FSC), the FSC cut-off criterion of 0.143 ^38^ was used.

### Production and purification of AAV2_L336C_ virus-like particles

The AAV2_L336C_ substitution was created within the AAV2 *cap* gene encoding all three viral proteins, VP1, VP2, and VP3, as previously described^4^. A recombinant baculovirus, encoding the AAV2 *cap* gene, with the L336C substitution, was created using the Bac-to-Bac system (Thermo Fisher). A plaque purified and titered baculovirus stock was used to infect *Sf*9 insect cells, at a multiplicity of infectivity of 5 to generate virus-like particles (VLPs). The harvested pellet (from lysed cells and polyethylene glycol precipitated supernatant) was freeze/thawed three times with Benzonase (EMD Millipore Cat#712053) treatment. After the third thaw, the resulting clarified supernatant was purified using a step iodixanol gradient followed by anion exchange^39^ and then dialyzed into 50 mM HEPES, pH 7.4 with 2 mM MgCl_2_, 150 mM NaCl. The sample concentration was determined by optical density assuming an extinction coefficient of 1.7 mg/(mL-cm) for AAV2 VLPs. The VLP purity and integrity were confirmed by sodium dodecyl sulfate polyacrylamide gel electrophoresis and negative stain EM on an FEI Spirit TEM, respectively.

### Single-Particle CryoEM Vitrification and Data Collection

Double blotting was used to increase particle concentration^40^. 2.5 μl of AAV2_L336C_ sample at 2.5 mg/ml was added to a plasma-cleaned (Gatan Solarus) 1.2 μm hole, 1.3 μm spacing holey gold grid (Quantifoil UltrAuFoil) and blotted away using Whatman grade 4 filter paper after 20s wait time. 2.5 μl of the same sample was then re-applied to the grid and blotted after 20s wait time and then vitrified in liquid ethane using a manual plunger. All operations were performed in a 4°C cold room at >80% humidity to minimize evaporation and sample degradation.

### Data Acquisition

Images were recorded on a Titan Krios electron microscope (FEI) equipped with a K2 summit direct detector (Gatan) at 0.394 Å per pixel in super-resolution counting mode (0.788 Å for the physical pixel size) using the Leginon software package^34^. Data collection was performed using a dose of ~22.5 e^−^/Å^2^ across 70 frames (50 msec per frame) at a dose rate of ~4.0 e^−^/pix/sec, using a set defocus range of −0.6 μm to −2 μm. On our microscope, the 100 μm objective aperture would allow for transmission of information up to ~1.4 Å, but could not be aligned to produce a coma-free diffractogram. In contrast, the 70 Åm aperture would truncate information at the ~2 Å limit. For this reason, the objective aperture was removed to prevent physical truncation of the most widely scattering electrons - and thus the highest-resolution information. Removal of the objective aperture in this case has the benefit of eliminating this aperture as a potential source of image astigmatism. A total of 1,317 micrographs were recorded over a single 3.5-day collection.

### Data Processing

Movie frames were aligned using MotionCor2^41^ with 5 by 5 patches, a grouping of 3 and B-factor of 100, and Fourier space binning of 2 (resulting in a pixel size of 0.788 Å/pixel) through the Appion software package^36^. Micrograph CTF estimation was performed using both CTFFind4^42^ for whole micrographs and GCTF^13^ for individual particles within the Appion software package. A subset of 8 micrographs was first used for particle picking using Gautomatch (Kai Zhang, unpublished), and particles were extracted and analyzed by 2D classification in Relion 2.1^11^. 2D class averages that showed clear structural details were used as templates for template-based picking using Gautomatch on all 1,317 micrographs. A total of 78,194 particles were then extracted using a box size of 800 pixels and subjected to two initial rounds of 2D classification (binned by 4) to identify and discard false positives such as ice and other obvious contaminants. Following 2D classification, 36,620 particles were reextracted with the re-centering option in Relion.

3D refinements were performed first using Relion and finishing in *cis*TEM^14^, with the initial model generated by CryoSPARC^43^. Icosahedral symmetry was imposed during all 3D refinement steps, based on prior knowledge^5^ of AAV2 structure. All conversions between Relion, CryoSPARC, and *cis*TEM were performed using Daniel Asarnow’s pyem script (unpublished). An initial 3D refinement using 7 rounds of auto-refinement and 2 rounds of local refinement with binning of 2. Particles were discarded based on analysis of the “score” values in *cis*TEM leading to the removal of a distinct subset of particles with low scores (below 6 in this dataset). This resulted in 30,515 particles that were re-extracted, unbinned and used for all subsequent operations. Per-particle CTF refinements were performed within *cis*TEM. All final refinements used a ring-shaped mask with an inner diameter of 75 Å and an outer diameter of 150 Å to specifically include only the capsid density and exclude remaining solvent. For this dataset, and after applying the stack-filtering procedures described above, 3D classifications did not produce any noticeable further gains.

Plotting the defocusV against defocusU values^44^ showed a systematic scaling of the difference between these two values as a function of their magnitude. Using the mag_distortion_estimate software^15^ and micrographs collected from a gold-coated cross grating replica grid (Supplementary Fig. 1), a magnification anisotropy of 1.10% was calculated. The appropriate correction for magnification anisotropy was applied during frame alignment (see above). The particle stack that was re-extracted from magnification-anisotropy-corrected frame sums reached a resolution of 1.97 Å after derivation of an *ab initio* model in CryoSparc and refinement in cisTEM.

The aligned movie frame stack was also split into individual frames and using the best Euler angles and shifts from above, reconstructions were computed using Frealign9. Frames 5–19, each of which independently exceeded a resolution of 2.24 Å, were summed and used for subsequent manual refinement (including CTF refinement) within *cis*TEM to obtain a reconstruction at 1.93 Å. The final reconstruction at 1.84 Å was computed after correcting for the curvature of the Ewald sphere using Frealign9. Rotational motion correction was performed in *cis*TEM by splitting each particle sum into groups of 5 frames (frames 5–9, 1014, and 15–19), and refining each group-of-5 as if it were a single particle. Particle-frame-averages with score lower than 3 were removed, resulting in a final stack of 87,781 “particles” that refined to 1.86 Å resolution. Although the nominal resolution at 0.143 cut-off was worse than without rotational frame alignment, inspection of the FSC curves and visual comparison of the two maps suggested that this procedure provided minor benefits. Notably, information at lower spatial frequencies was slightly improved within the reconstruction following rotational frame alignment.

To generate maps of the opposite handedness, Euler angles of the particles were changed from (phi, theta, psi) to (-phi, 180-theta, psi). Ewald sphere curvature corrected reconstructions of same and opposite handedness were done by setting IEWALD to either 1 or −1 respectively in Frealign9.

### AAV2_L336C_ model refinement

For model refinement of the AAV2_L336C_ variant, the deposited structure of AAV2 (PDB-ID: 1LP3) was used as a starting template. A 60mer capsid model downloaded from VIPERdb^45^ (http://viperdb.scripps.edu) was docked into the map using the ‘Fit-in-map’ function in the Chimera^46^ program. To optimize the correlation coefficient (CC) between the model and map the voxel (pixel) size of the map was adjusted. From the fitted 60mer, a monomer was extracted for the model building. For model building and real space refinement in the Coot^47^ program, the map was converted from the Purdue Image Format (PIF) to the XPlor format using e2proc3D.py subroutine in the EMAN2^48^ application and finally to the CCP4 format using MAPMAN^49^. In Coot^47^ L336 in AAV2_WT_ was substituted to a cysteine and the side-chain and main-chain atoms, including those of neighboring residues, adjusted to better fit the experimental density map using the real-space-refinement subroutine. After the manual refinement of the monomer was completed, a 60mer was regenerated in VIPERdb by T=1 icosahedral matrix multiplication and the model refined against the cryo-EM map utilizing the real space, and B-factor refinement subroutines in the Phenix^24^ program. The CC and refinement statistics, including root mean square deviations (RMSD), bond lengths and angles were analyzed by Phenix. Model adjustment and refinement were performed iteratively in Coot and Phenix, and the statistics were examined using Molprobity^50^ until no further improvements were observed. The final map and model were then validated using 1) EMRinger^23^ to compare map to model, 2) SPARX^28^ to calculate map local resolution and 3) 3DFSC program suite^27^ to calculate degree of directional resolution anisotropy through the 3DFSC.

## References for Main Text

1. Elmlund, D., Le, S.N. & Elmlund, H. High-resolution cryo-EM: the nuts and bolts. Current opinion in structural biology 46, 1-6 (2017).

2. Merk, A. et al. Breaking cryo-EM resolution barriers to facilitate drug discovery. Cell 165, 1698-1707 (2016).

3. Bleker, S., Sonntag, F. & Kleinschmidt, J.A. Mutational analysis of narrow pores at the fivefold symmetry axes of adeno-associated virus type 2 capsids reveals a dual role in genome packaging and activation of phospholipase A2 activity. Journal of virology 79, 2528-2540 (2005).

4. Grieger, J.C., Johnson, J.S., Gurda-Whitaker, B., Agbandje-McKenna, M. & Samulski, R.J. Surface-exposed adeno-associated virus Vp1-NLS capsid fusion protein rescues infectivity of noninfectious wild-type Vp2/Vp3 and Vp3-only capsids but not that of fivefold pore mutant virions. Journal of virology 81, 7833-7843 (2007).

5. Agbandje-McKenna, M. & Kleinschmidt, J. in Adeno-Associated Virus 47-92 (Springer, 2012).

6. Lu, Y. Recombinant adeno-associated virus as delivery vector for gene therapy—a review. Stem cells and development 13, 133-145 (2004).

7. Snijder, J. et al. Defining the stoichiometry and cargo load of viral and bacterial nanoparticles by Orbitrap mass spectrometry. Journal of the American Chemical Society 136, 7295-7299 (2014).

8. Danev, R. & Baumeister, W. Cryo-EM single particle analysis with the Volta phase plate. Elife 5 (2016).

9. Fischer, N. et al. Structure of the E. coli ribosome-EF-Tu complex at< 3 Å resolution by C s-corrected cryo-EM. Nature 520, 567 (2015).

10. Campbell, M.G., Veesler, D., Cheng, A., Potter, C.S. & Carragher, B. 2.8 Å resolution reconstruction of the Thermoplasma acidophilum 20S proteasome using cryo-electron microscopy. Elife 4 (2015).

11. Scheres, S.H. RELION: implementation of a Bayesian approach to cryo-EM structure determination. J Struct Biol 180, 519-530 (2012).

12. Grigorieff, N. Frealign: An Exploratory Tool for Single-Particle Cryo-EM. Methods Enzymol 579, 191-226 (2016).

13. Zhang, K. Gctf: Real-time CTF determination and correction. J Struct Biol 193, 1-12 (2016).

14. Grant, T., Rohou, A. & Grigorieff, N. cisTEM, user-friendly software for single-particle image processing. Elife 7 (2018).

15. Grant, T. & Grigorieff, N. Automatic estimation and correction of anisotropic magnification distortion in electron microscopes. Journal of Structural Biology 192, 204-208 (2015).

16. Wolf, M., DeRosier, D.J. & Grigorieff, N. Ewald sphere correction for single-particle electron microscopy. Ultramicroscopy 106, 376-382 (2006).

17. Liu, Z., Guo, F., Wang, F., Li, T.-C. & Jiang, W. 2.9 Å resolution cryo-EM 3D reconstruction of close-packed virus particles. Structure 24, 319-328 (2016).

18. Crowther, R. Procedures for three-dimensional reconstruction of spherical viruses by Fourier synthesis from electron micrographs. Phil. Trans. R. Soc. Lond. B 261, 221-230 (1971).

19. DeRosier, D.J. Correction of high-resolution data for curvature of the Ewald sphere. Ultramicroscopy 81, 83-98 (2000).

20. Jensen, G.J. & Kornberg, R.D. Defocus-gradient corrected back-projection. Ultramicroscopy 84, 57-64 (2000).

21. Zhang, X. & Zhou, Z.H. Limiting factors in atomic resolution cryo electron microscopy: no simple tricks. Journal of structural biology 175, 253-263 (2011).

22. Russo, C.J. & Henderson, R. Ewald sphere correction using a single side-band image processing algorithm. Ultramicroscopy (2018).

23. Barad, B.A. et al. EMRinger: side chain-directed model and map validation for 3D cryo-electron microscopy. Nature methods 12, 943 (2015).

24. Adams, P.D. et al. PHENIX: a comprehensive Python-based system for macromolecular structure solution. Acta Crystallographica Section D: Biological Crystallography 66, 213-221 (2010).

25. Herzik Jr, M.A., Wu, M. & Lander, G.C. Achieving better-than-3-A resolution by singleparticle cryo-EM at 200 keV. nature methods 14, 1075 (2017).

26. Stagg, S.M., Noble, A.J., Spilman, M. & Chapman, M.S. ResLog plots as an empirical metric of the quality of cryo-EM reconstructions. Journal of structural biology 185, 418-426 (2014).

27. Tan, Y.Z. et al. Addressing preferred specimen orientation in single-particle cryo-EM through tilting. Nature methods 14, 793 (2017).

28. Hohn, M. et al. SPARX, a new environment for Cryo-EM image processing. Journal of structural biology 157, 47-55 (2007).

## Online Methods References

29. Russo C.J. & Passmore L.A. Ultrastable gold substrates for electron cryomicroscopy. Science 346, 1377-1380 (2014).

30. Glaeser, R.M., Typke, D., Tiemeijer, P.C., Pulokas, J. & Cheng, A. Precise beam-tilt alignment and collimation are required to minimize the phase error associated with coma in highresolution cryo-EM. Journal of structural biology 174, 1-10 (2011).

31. Li X. et al. Electron counting and beam-induced motion correction enable near-atomic-resolution single-particle cryo-EM. Nature methods 10, 584 (2013).

32. McMullan, G., Faruqi, A., Clare, D. & Henderson, R. Comparison of optimal performance at 300 keV of three direct electron detectors for use in low dose electron microscopy. Ultramicroscopy 147, 156-163 (2014).

33. Brilot A.F. et al. Beam-induced motion of vitrified specimen on holey carbon film. Journal of structural biology 177, 630-637 (2012).

34. Suloway C. et al. Automated molecular microscopy: the new Leginon system. Journal of structural biology 151, 41-60 (2005).

35. Xie Q. et al. The atomic structure of adeno-associated virus (AAV-2), a vector for human gene therapy. Proceedings of the National Academy of Sciences 99, 10405-10410 (2002).

36. Lander G.C. et al. Appion: an integrated, database-driven pipeline to facilitate EM image processing. J Struct Biol 166, 95-102 (2009).

37. Passos D.O. & Lyumkis D. Single-particle cryoEM analysis at near-atomic resolution from several thousand asymmetric subunits. Journal of structural biology 192, 235-244 (2015).

38. Rosenthal P.B. & Henderson R. Optimal determination of particle orientation, absolute hand, and contrast loss in single-particle electron cryomicroscopy. J Mol Biol 333, 721-745 (2003).

39. Drouin L.M. et al. Cryo-electron microscopy reconstruction and stability studies of the wild type and the R432A variant of adeno-associated virus type 2 reveal that capsid structural stability is a major factor in genome packaging. Journal of virology 90, 8542-8551 (2016).

40. Snijder J. et al. Vitrification after multiple rounds of sample application and blotting improves particle density on cryo-electron microscopy grids. Journal of structural biology 198, 38-42 (2017).

41. Zheng S.Q. et al. MotionCor2: anisotropic correction of beam-induced motion for improved cryo-electron microscopy. Nat Methods 14, 331-332 (2017).

42. Rohou A. & Grigorieff N. CTFFIND4: Fast and accurate defocus estimation from electron micrographs. J Struct Biol 192, 216-221 (2015).

43. Punjani, A., Rubinstein, J.L., Fleet, D.J. & Brubaker, M.A. cryoSPARC: algorithms for rapid unsupervised cryo-EM structure determination. Nat Methods 14, 290-296 (2017).

44. Zhao, J., Brubaker, M.A., Benlekbir, S. & Rubinstein, J.L. Description and comparison of algorithms for correcting anisotropic magnification in cryo-EM images. Journal of structural biology 192, 209-215 (2015).

45. Shepherd C.M. et al. VIPERdb: a relational database for structural virology. Nucleic acids research 34, D386-D389 (2006).

46. Pettersen E.F. et al. UCSF Chimera—a visualization system for exploratory research and analysis. Journal of computational chemistry 25, 1605-1612 (2004).

47. Emsley P. & Cowtan K. Coot: model-building tools for molecular graphics. Acta Crystallographica Section D: Biological Crystallography 60, 2126-2132 (2004).

48. Tang G. et al. EMAN2: an extensible image processing suite for electron microscopy. Journal of structural biology 157, 38-46 (2007).

49. Kleywegt G.J. & Jones T.A. xdlMAPMAN and xdlDATAMAN-programs for reformatting, analysis and manipulation of biomacromolecular electron-density maps and reflection data sets. Acta Crystallographica Section D: Biological Crystallography 52, 826-828 (1996).

50. Chen V.B. et al. MolProbity: all-atom structure validation for macromolecular crystallography. Acta Crystallographica Section D: Biological Crystallography 66, 12-21 (2010).

